# Sympathetic activation and the force-frequency relationship in heart failure with reduced ejection fraction

**DOI:** 10.64898/2026.08.24.746885

**Authors:** Sam Straw, Ankit Gupta, Beatrice Bretherton, Charlotte Cole, Oliver I Brown, Stephe Kamalathasan, Michael Drozd, Judith Lowry, Julie Corrigan, Maria F Paton, Ruth Burgess, Mark T Kearney, Richard M Cubbon, Klaus K Witte, John Gierula

**Affiliations:** Leeds Institute of Cardiovascular and Metabolic Medicine, University of Leeds, Leeds, UK; Leeds Teaching Hospitals NHS Trust, UK; School of Biomedical Sciences, University of Leeds, Leeds, UK

**Author notes:** denotes joint first authorship (Sam Straw, Ankit Gupta). **Corresponding author**: Dr Klaus K Witte. Leeds Institute of Cardiovascular and Metabolic Medicine, LIGHT Building, University of Leeds, Clarendon Way, Leeds, LS2 9JT, UK.

**Keywords:** Heart failure, heart rate, sympathetic, autonomic, haemodynamic

## Abstract

**Background:** Limited heart rate rise contributes to reduced exercise tolerance for people who have heart failure with reduced ejection fraction (HFrEF), yet rate-adaptive pacing does not improve functional capacity due to an attenuated force-frequency relationship (FFR). How the FFR relates to total peripheral resistance and sympathetic tone in HFrEF is unknown.

**Methods:** In a prospective, observational study, participants with HFrEF and controls underwent an incremental pacing protocol, during which heart rate was increased from 50 to 140 beats per minute. At each heart rate increment LV contractility was measured by echocardiography to determine the FFR, as well as continuous beat-to-beat measurement of systolic and diastolic blood pressures with a plethysmography device to determine cardiac output, total peripheral resistance and blood pressure variability (BPV). A microneurography study was then conducted to measure muscle sympathetic nerve activity (MSNA) during incremental pacing.

**Results:** A total of 157 participants with HFrEF and 55 controls (mean age 71.1±1.4 years, 172 (81.1%) male) underwent the pacing protocol. We observed single units in seven of 11 participants who participated in the microneurography study. In both groups, LV contractility and cardiac output increased until the peak of the FFR, after which these declined. We observed a reduction in total peripheral resistance, blood pressure variability, MSNA frequency and incidence coinciding with the peak of the FFR, beyond which these increased. Whilst these relationships were present in both groups, they were more evident in participants with HFrEF.

**Conclusions:** For people with HFrEF there is a bidirectional relationship between heart rate and sympathetic activation, with a nadir of sympathetic tone occurring at the peak of the FFR. Both excessively low and high heart rates are accompanied by greater sympathetic activation. Taken together, these data suggest that optimal heart rate targets for HFrEF are likely to be individual.

## Introduction

### Background

Heart rate is a predictor of cardiovascular risk in the general population^1^ and for people with established cardiovascular diseases including heart failure (HF).^2,3^ For individuals who have heart failure with reduced ejection fraction (HFrEF), every 5 beats per minute (bpm) increase in resting heart rate is associated with 16% greater risk of cardiovascular mortality or HF hospitalisation.^4^ Treatments which lower heart rate, including beta-adrenoceptor antagonists, improve left ventricular (LV) function^5^ and survival,^6^ whilst selective *I*_f_ channel inhibition reduces the risk of HF hospitalisation.^7^ Despite their beneficial effects on clinical outcomes, limited heart rate rise during physical activity commonly contributes to reduced exercise tolerance for individuals receiving these agents, as cardiac output is a function of heart rate and stroke volume.^8^ However, imprecise increases in heart rate using rate-adaptive pacing consistently fail to show benefits on functional capacity in HF, possibly because the relationship between heart rate and LV contractility (the force-frequency relationship [FFR]) is not linear.^9,10^

It was previously shown that when heart rate increases during exercise are guided by the FFR, walk distance and LV function are improved after 6-months.^11^ However, the acute physiological consequences of changes to heart rate and how these relate to the FFR, total peripheral resistance and sympathetic activation in HF have not been defined. We hypothesised that increasing heart rate with cardiac pacing would, through increased cardiac output and baroreceptor loading result in the withdrawal of sympathetic activation. Using an incremental cardiac pacing protocol with simultaneous cardiac imaging, non-invasive continuous beat-to-beat haemodynamic assessment, and microneurography, it is demonstrated for the first time how the FFR is related to sympathetic activation. Given sympathetic activation is itself a driver of increased resting heart rates, our findings suggest the FFR underpins a homeostatic loop coupling heart rate with sympathetic tone.

## Methods

### Study design

A prospective, observational, mechanistic study was conducted, enrolling participants who had previously received cardiac implantable electronic devices (CIED) according to standard indications.^12^ We compared people who had CIEDs implanted for the management of HFrEF with participants without heart failure, who had previously received CIEDs for the management of bradycardia and served as controls. The United Kingdom Health Research Authority provided ethical approval for the study (Yorkshire & The Humber – Leeds West Research Ethics Committee 20/YH/0087). All participants provided written, informed consent and the study complied with the principles outlined in the Declaration of Helsinki. The study protocol is registered with Clinical Trials (NCT04201015).

### Study participants

Participants with HFrEF were eligible if they had symptoms of HF, left ventricular ejection fraction (LVEF) <50%, existing CIEDs implanted according to standard indications, and were receiving stable doses of medications for HF for at least three months. For controls, we required participants to not have current signs or symptoms or prior diagnosis of HF, or any prior measurement of LVEF <50%. We also excluded any individual who fulfilled the diagnostic criteria for heart failure with preserved ejection fraction.^13^ For all participants we recorded demographic details, past medical history and currently prescribed medications. We recorded height and weight, and calculated body surface area using the Mosteller equation.^14^

### Experimental protocol

At the beginning of the study visit, CIEDs were interrogated to record programmed mode and parameters, generator voltage, lead capture thresholds, sensing and impedances. Throughout, CIED interrogation and programming were done using wireless telemetry where possible, or with the telemetry header positioned over the device to allow for reprogramming during the study protocol. Assessments were made from a programmed base rate of 50bpm, or at the participant’s intrinsic resting heart rate where this was >50bpm. The programmed base rate was then increased by 15bpm increments at each stage up to a maximum of 140bpm. Where participants did not have biventricular pacemakers or defibrillators, we minimised right ventricular pacing by either programming an atrial pacing mode or extending atrioventricular delays to allow for intrinsic atrioventricular conduction. At each heart rate we recorded systolic and diastolic blood pressure measured using a standard sphygmomanometer placed on the participant’s right arm, recorded at each heart rate increment at the point where the first tapping sound occurred for two consecutive beats. We also recorded continuous beat-to-beat systolic and diastolic blood pressures and cardiac output using plethysmography. Cardiac imaging was done at each heart rate according to a standardised imaging acquisition protocol. Following the experimental protocol, a further device interrogation was done, and devices were programmed to their original settings. Participants were then observed for at least 30 minutes.

### Cardiac imaging acquisition protocol and analysis

Participants were imaged in a left lateral position using a cardiac ultrasound imaging system (Vivid 95 Cardiac Ultrasound, GE Healthcare, Chicago, USA) by a cardiac sonographer accredited by the British Society of Echocardiography. We acquired images to obtain a minimum echocardiographic dataset, recorded at the participant’s baseline heart rate. During the experimental protocol we acquired apical two and four-chamber images at each heart rate increment. Images were sent to digital storage media and, for the present analysis were analysed offline using Medcon (McKesson Cardiology, Irving TX, USA). Where endocardial border definition allowed, LV end-diastolic and end-systolic volumes were measured using the biplane method of disks, indexed for body surface area.

### Determination of the force frequency relationship and critical heart rate

The end-systolic pressure-volume relationship represents an afterload-independent assessment of LV contractility.^15^ We used systolic blood pressure measured with a manual sphygmomanometer as a surrogate of LV end-systolic pressure and determined LV end-systolic volume using transthoracic echocardiography, as previously described.^16,17^ LV contractility was expressed as the ratio of systolic blood pressure to LV end-systolic volume indexed for body surface area.^18^ LV contractility was plotted against heart rate to describe the force-frequency relationship for each individual participant. We defined the ‘critical’ heart rate as the heart rate coinciding with the peak of the force-frequency relationship curve.

### Continuous beat-to-beat non-invasive haemodynamic assessment

Participants lay recumbent in a left lateral position having rested supine for at least five minutes. Continuous beat-to-beat measurement of systolic and diastolic blood pressures were made non-invasively using a plethysmography device (Finometer, Finapres Medical System, Netherlands). This device uses a cuff placed around the index finger, with a photoelectric plethysmograph and a volume-clamp circuit to provide a continuous beat-to-beat arterial blood pressure waveform. This device also determined cardiac output and total peripheral resistance and blood pressure at each heart rate. Heart rate during this assessment was recorded by a three-lead electrocardiogram with surface electrodes placed on the clavicles and the costal margins. Given that CIEDs deliver digitally timed pacing rates, determination of heart rate variability was not possible. We therefore determined beat-to-beat variability of systolic blood pressure, as a surrogate marker of sympathetic tone. Blood pressure variability (BPV) was expressed firstly by the coefficient of variation (CoV), calculated as the ratio between the standard deviation and the mean, and secondly as the average real variability (ARV), calculated by averaging the absolute difference between consecutive beat-to-beat systolic blood pressure readings.

### Microneurography

We undertook an exploratory microneurography study to confirm our non-invasive findings, recording muscle sympathetic nerve activity (MSNA) in 11 participants with HFrEF. For this study participants lay semi-supine and the common peroneal nerve was identified by manual palpation below the head of the fibula, as previously described.^19^ A tungsten microelectrode (FHC Inc, Bowdoin, ME) was inserted percutaneously into the peroneal nerve (recording electrode) with a second into the subcutaneous tissue approximately 1-2cm away (reference electrode). The nerve signal was amplified (x50k), filtered and digitized. The data were displayed in real time using Spike2 (Cambridge Electronic Design Limited, Cambridge, UK) allowing the inspection of the nerve signal during the study procedures with the recording electrode manipulated until a single unit could be visualized. During the recording we used contralateral isometric handgrip exercise consisting of squeezing a dynamometer at ∼50% perceived voluntary contraction for 2 minutes.

Nerve and blood pressure traces were exported from LabChart 8 (AD Instruments, Sydney, Australia) in a single file and imported into Spike2 (Cambridge Electronic Design, Milton, UK). We required single units to have occurred in diastole, increased in firing with increases in blood pressure, have a higher firing rate during the second half than during the first half of an isometric handgrip test, and required the shape and rate of the units to not change during the stroking of the upper foot. Units which did not meet any of these criteria were discarded. Further confirmation was obtained during offline analysis (CorelDRAW, version 6, Ottawa, Canada) by superimposing all putative MSNA units to ensure the amplitude and shape remained constant indicating these were recorded from the same axon (Figure S1).

MSNA recordings were made from a programmed base rate of 50bpm, or at the participant’s intrinsic resting heart rate where this was >50bpm. The programmed base rate was then increased by 15bpm increments, with a recording of at least three minutes made for each stage up to a maximum of 140bpm. Single units were counted manually for each heart rate recording. MSNA frequency (per minute) was calculated by counting all single units that occurred in the recording period divided by the duration of the recording period. MSNA incidence (number of units per 100 heart beats) was also derived to account for the effect of any changes in heart rate not related to true increases in MSNA activity, by dividing MSNA frequency by the mean heart rate and multiplying by 100.

### Statistical analysis

Continuous data are presented as means ± standard deviation, or median (interquartile range) unless otherwise stated. Categorical data are presented as counts and percentages. After testing for normality, differences between groups were assessed using Student’s t-test for continuous data, and by chi-squared test for categorical data. We present haemodynamic data as mean values for each heart rate with their standard error, and also as percentage changes relative to each individual’s critical heart rate, compared by two-way analysis of variance. Statistical analyses were done using R version 4.1.1 (R Core Team) with figures illustrated using Prism version 10.6.1 (GraphPad Software Inc).

## Results

### Participant characteristics

Between 1^st^ November 2022 and 27^th^ June 2024, we recruited a total of 217 participants who underwent echocardiographic and plethysmographic assessment during an incremental pacing protocol. Following the exclusion of 5 participants who fulfilled diagnostic criteria for heart failure with preserved ejection fraction, our dataset comprised 157 participants with HFrEF and 55 controls. The mean age was 71.1±1.4 years and 172 (81.1%) were male. Of these, a total of 210 had sufficient endocardial border definition to assess LV end-systolic and end-diastolic volumes. Eleven participants with HFrEF subsequently underwent the microneurography study. Data contrasting participants with HFrEF and controls are displayed in Table 1. Participants with HFrEF were similar in age, sex distribution and body mass index compared to controls, but were more likely to have history of ischaemic heart disease and had on average higher serum creatinine. For those with HFrEF, the mean LVEF was 42.5±8.0% and the mean NT-proBNP was 969.5 ± 1221.6pg/L. Of those with HFrEF, 110 (70.1%) were receiving an angiotensin converting enzyme inhibitor, angiotensin receptor blocker, or angiotensin receptor-neprilysin inhibitor, 120 (76.4%) a beta-adrenoceptor antagonist, 65 (41.4%) a mineralocorticoid receptor antagonist, and 53 (33.7%) were prescribed a loop diuretic.

**Table 1.** Demographics and clinical characteristics of participants comparing those with HFrEF and controls.

|  | **LVEF ≤ 50%**<br>(n = 157) | **LVEF > 50%**<br>(n = 55) | **p-value** |
| --- | --- | --- | --- |
| **Participant demographics** |  |  |  |
| **Male sex, n (%)** | 131 (83.4) | 41 (74.5) | 0.147 |
| **Age, years** | 72.1 (10.5) | 70.7 (11.0) | 0.401 |
| **Height, cm** | 173.3 (7.9) | 170.4 (8.4) | **0.025** |
| **Weight, kg** | 84.6 (15.7) | 79.8 (15.0) | 0.053 |
| **Body mass index, kg/m<sup>2</sup>** | 48.7 (8.2) | 46.7 (7.8) | 0.123 |
| **Clinical history** |  |  |  |
| **Atrial fibrillation, n (%)** | 60 (38.2) | 14 (25.5) | 0.082 |
| **Cerebrovascular disease, n (%)** | 14 (8.9) | 8 (14.5) | 0.302 |
| **Diabetes mellitus, n (%)** | 40 (25.5) | 9 (16.4) | 0.161 |
| **Hypertension, n (%)** | 66 (42.0) | 30 (54.5) | 0.092 |
| **Ischaemic heart disease, n (%)** | 62 (39.5) | 10 (18.2) | **0.005** |
| **Medications** |  |  |  |
| **ACE inhibitor/ARB/ARNI, n (%)** | 110 (70.1) | 18 (32.7) | **< 0.001** |
| **Aspirin, n (%)** | 29 (18.5) | 10 (18.2) | 0.900 |
| **Beta-blocker, n (%)** | 120 (76.4) | 21 (38.2) | **< 0.001** |
| **Calcium channel blocker, n (%)** | 24 (15.3) | 13 (23.6) | 0.173 |
| **Loop diuretic, n (%)** | 53 (33.7) | 5 (9.1) | **< 0.001** |
| **MRA, n (%)** | 65 (41.4) | 0 (0.0) | **< 0.001** |
| **SGLT2 inhibitor, n (%)** | 34 (21.7) | 0 (0.0) | **< 0.001** |
| **Statin, n (%)** | 105 (66.9) | 38 (69.1) | 0.854 |

| Blood tests |  |  |  |
| --- | --- | --- | --- |
| NT-pro BNP, pg/ml | 969.5 (1221.6) | 466.0 (852.3) | **0.009** |
| Haemoglobin, g/l | 143.6 (18.6) | 146.4 (11.0) | 0.393 |
| Creatinine, mmol/L | 103.5 (35.1) | 86.8 (20.5) | **0.005** |
| Echocardiography |  |  |  |
| LVEDVi, ml/m<sup>2</sup> | 66.0 (25.0) | 52.2 (13.6) | **< 0.001** |
| LVESVi, ml/m<sup>2</sup> | 38.6 (18.3) | 22.9 (7.3) | **< 0.001** |
| LVEF, % | 42.5 (8.0) | 57.5 (5.7) | **< 0.001** |
| LV contractility, mmHg/ml/m<sup>2</sup> | 2.5 (1.1) | 2.7 (0.8) | 0.248 |

LVEF; left ventricular ejection fraction, ACE; angiotensin converting enzyme, ARB; angiotensin receptor blocker, ARNI; angiotensin-receptor neprilysin inhibitor, MRA; mineralocorticoid receptor antagonist, SGLT2; sodium glucose co-transporter, NT-proBNP; N-terminal B-type natriuretic peptide, LVEDVi; indexed left ventricular end-diastolic volume, LVESVi; indexed left ventricular end-systolic volume, LV; left ventricular.

### Relationship between heart rate, indices of left ventricular contractility and cardiac output

We determined the FFR for each individual non-invasively by increasing heart rate using an incremental pacing protocol and measuring systolic blood pressure and LV end-systolic volume by echocardiography at each heart rate. For both participants with HFrEF and for controls, we observed increases in LV contractility as heart rate increased, until a critical heart rate, beyond which it declined (Figures 1A and 1B). LV contractility was, on average, lower in participants with HFrEF than in controls (*p*<0.001), and there was a significant interaction between participants with HFrEF and controls with respect to the relationship between LV contractility and heart rate (p=0.04), whereby the slope of the force-frequency relationship was less in participants with HFrEF. Relative to LV contractility at the critical heart rate, LV contractility was lower for all heart rates for both groups (*p*=0.006), however the relative changes were similar. We observed a similar relationship between heart rate and the rate of pressure change (dP/dt) determined by non-invasive plethysmography, although the reduction in dP/dt beyond the critical heart rate was more evident in those with HFrEF than in controls (Figures 1C and 1D). Cardiac output was similar comparing those with HFrEF and controls (*p*=0.22) (Figures 1E and 1F). In both groups cardiac output was lower for heart rate above, and below the critical heart rate (*p*<0.001).

**Figure 1.**
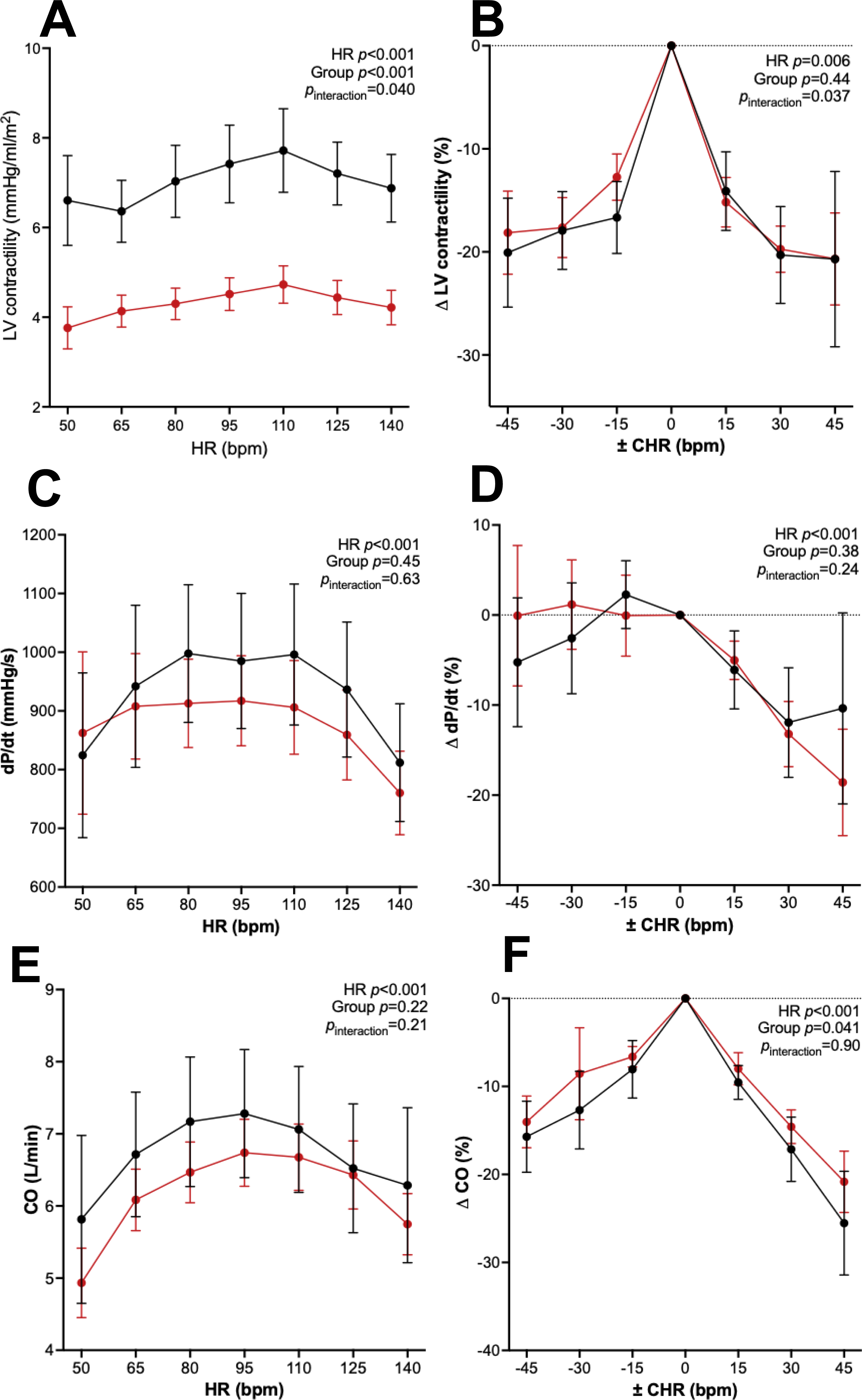
Relationship between heart rate, indices of LV contractility, and cardiac output. <u>Legend</u> Plots demonstrating the relationship between **A** LV contractility and heart rate, **B** relative change in LV contractility compared to the critical heart rate, **C** rate of pressure change and heart rate, **D** relative change in rate of pressure change compared to the critical heart rate, **E** cardiac output and heart rate, and **F** relative change in cardiac output compared to the critical heart rate, comparing participants with HFrEF and controls.

### Relationship amongst heart rate, systolic and diastolic blood pressure and mean arterial pressure

The relationship between heart rate and systolic blood pressure was negatively correlated and curvilinear and was similar comparing participants with HFrEF and controls (*p*=0.46). Systolic blood pressure was, on average higher in participants with HFrEF below the critical heart rate and then lower beyond this for both groups (*p*<0.001) (Figure 2 A and B). In contrast, diastolic blood pressure increased for both groups with increasing heart rate and was similar comparing participants with HFrEF and controls (*p*=0.81) (Figure 2 C and D). As a consequence of these divergent changes in systolic and diastolic blood pressure, mean arterial pressure increased until the critical heart rate, beyond which it remained similar (Figures 2 E and F).

**Figure 2.**
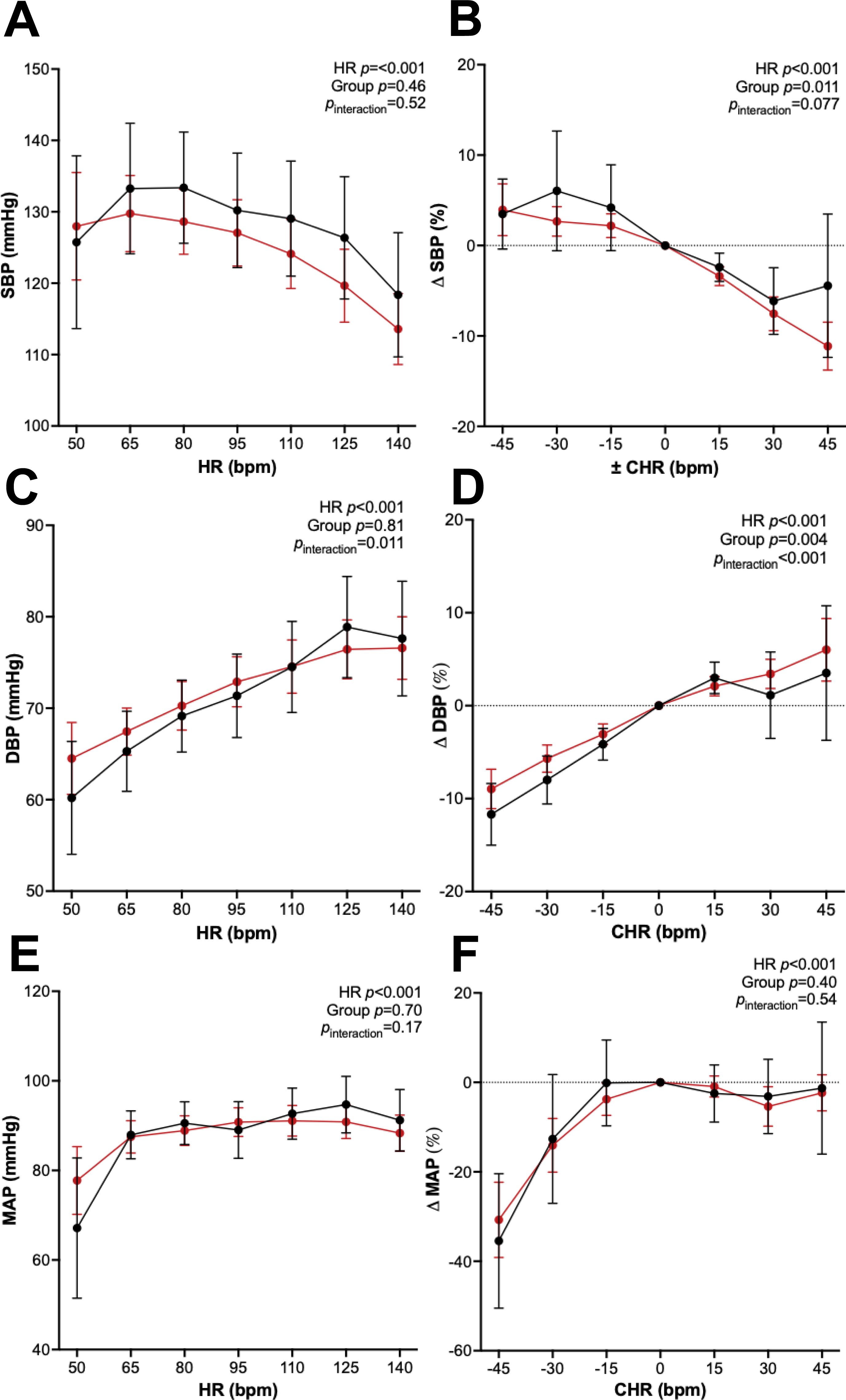
Relationship amongst heart rate, systolic and diastolic blood pressure and mean arterial pressure. <u>Legend</u> Plots demonstrating the relationship between **A** systolic blood pressure and heart rate, **B** relative change in systolic blood pressure compared to the critical heart rate, **C** diastolic blood pressure and heart rate, **D** relative change in diastolic blood pressure compared to the critical heart rate, **E** mean arterial pressure and heart rate, **F** relative change in mean arterial pressure compared to the critical heart rate, comparing participants with HFrEF and controls.

### Relationship between heart rate and blood pressure variability

An inverse relationship was observed between heart rate and BPV for both participants with HFrEF and controls, with a reduction in BPV around the critical heart rate (*p*=0.50) (Figures 3A-D). The relationship was more evident in participants with HFrEF, who had, on average, greater coefficient of variation at all heart rates below and above the critical heart rate. We also observed greater average real variability (ARV) for participants with HFrEF and controls at heart rates below and above the critical heart rate (*p*=0.003).

**Figure 3.**
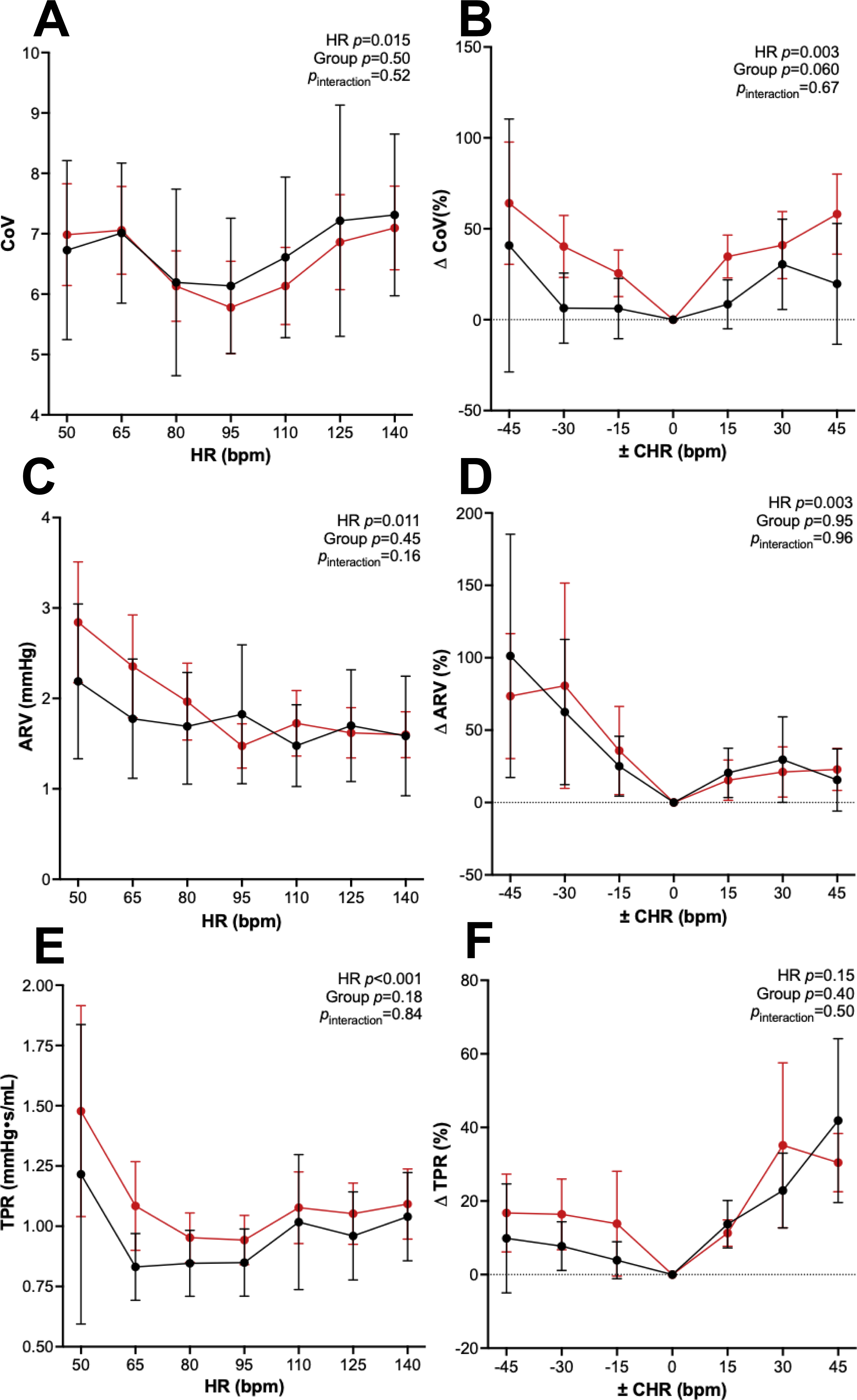
Relationship amongst heart rate, vascular resistance, and blood pressure variability. <u>Legend</u> Plots demonstrating the relationship between **A** total peripheral resistance and heart rate, and **B** relative change in total peripheral resistance compared to the critical heart rate, **C** coefficient of variation of systolic blood pressure and heart rate, **D** relative change in coefficient of variation of systolic blood pressure compared to the critical heart rate, **E** average real difference of beat-to-beat systolic blood pressure and heart rate, and **F** relative change in average real difference of beat-to-beat systolic blood pressure compared to the critical heart rate, comparing participants with HFrEF and controls.

### Relationship between heart rate and total peripheral resistance

This inverse relationship was also observed for heart rate and total peripheral resistance. In participants with HFrEF, total peripheral resistance reached its nadir at the critical heart rate. This relationship was also observed in controls, although was less evident below the critical heart rate at which total peripheral resistance was similar. We found that the greatest total peripheral resistance in absolute terms occurred at the lowest heart rates in participants with HFrEF.

### Relationship between heart rate and muscle sympathetic nerve activity

We supported these observations in an exploratory microneurography study in 11 participants with HFrEF. All 11 participants completed the study protocol, and we were able to confirm the presence of single units in seven. In these seven participants, the MSNA frequency relationship mirrored that of total vascular resistance and BPV, with a nadir of sympathetic tone with lower frequency and incidence of MSNA as heart rate increased, until the critical heart rate, beyond which it was greater (Figures 4A and 4B). When accounting for the increase in heart rate during the pacing protocol, we found a similar pattern with MSNA incidence, reducing as heart rate increased, and then increasing beyond the critical heart rate (Figures 4C and 4D).

**Figure 4.**
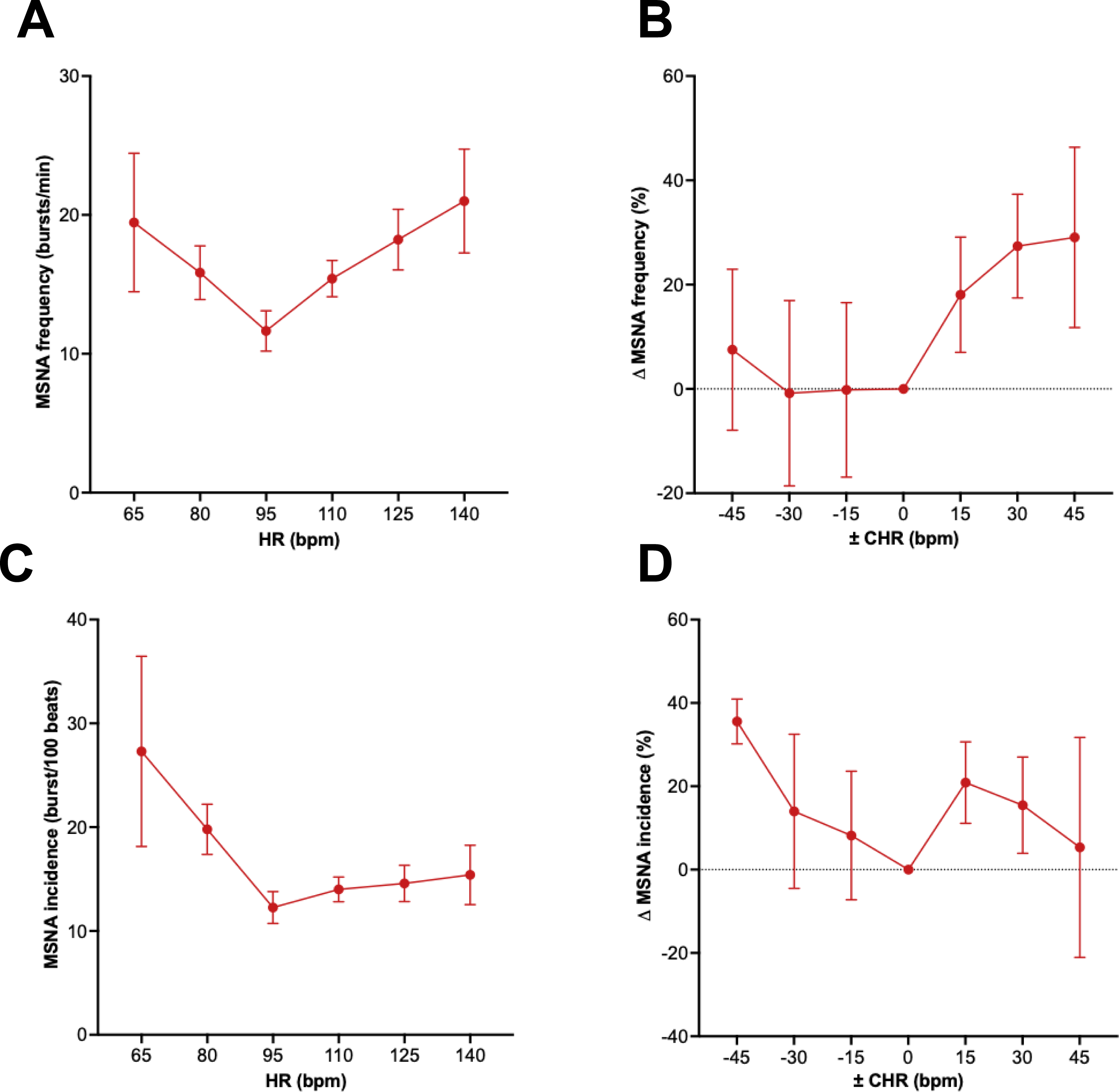
Relationship between heart rate and MSNA. <u>Legend</u> Plots demonstrating the relationship between **A** MSNA frequency and heart rate, **B** relative change in MSNA frequency compared to the critical heart rate, **C** MSNA incidence and heart rate, and **D** relative change in MSNA incidence compared to the critical heart rate, in participants with HFrEF.

## Discussion

### Principal findings

In this prospective, observational, mechanistic study we defined for each individual how heart rate related to LV contractility, cardiac output, beat-to-beat blood pressure, and total peripheral resistance. Using the FFR as a common reference, we found that the fall in vascular resistance coincided with the critical heart rate, accompanied by a reduction in directly measured MSNA. These observations suggest a coupling between the FFR and sympathetic outflow, and that both excessively high and excessively low heart rates are accompanied by greater sympathetic tone and vascular resistance. By comparing people with HFrEF and controls, we show that this bidirectional relationship between heart rate and sympathetic outflow is not disease specific, rather a general homeostatic mechanism, although this was exaggerated in HFrEF.

### Sympathetic activation in heart failure

The syndrome of HF results from a complex interplay between structural and functional changes of the LV, renin-angiotensin-aldosterone system, autonomic nervous system, peripheral vasculature, and skeletal muscle.^20^ Regardless of whether this follows an acute or chronic insult, the pathophysiology of HFrEF is similar and underpinned by LV dilatation, reduced contractile function and a fall in cardiac output insufficient to meet the body’s metabolic requirements. This initial decline in cardiac output is detected by arterial baroreceptors which then trigger a series of homeostatic reflexes initiated by an increase in sympathetic and withdrawal of parasympathetic tone.^21,22^ This baroreceptor-mediated increase in sympathetic tone is responsible for the initial overactivation of the renin-angiotensin-aldosterone system, and in the long term elevates resting heart rate increasing myocardial oxygen demand, causing vasoconstriction, driving adverse remodelling and increasing automaticity predisposing to ventricular tachyarrhythmias.

### Bidirectional coupling of the force-frequency relationship and sympathetic activation

The FFR describes the relationship between LV contractility and heart rate, with its peak defining a critical heart rate at which contractility is greatest. We hypothesised that increases in heart rate up until the peak of the FFR would, by augmenting cardiac output and arterial pressure, result in loading of arterial baroreceptors, causing vasodilatation and a fall in total peripheral resistance. Through non-invasive assessments using echocardiography and plethysmography, we observed that increases in heart rate were accompanied by increases in LV contractility, cardiac output, and mean arterial pressure up to the peak of the FFR and that total peripheral resistance was lowest at the critical heart rate. However, as total peripheral resistance is derived from the ratio of mean arterial pressure and cardiac output, this observation is not independent from the changes in LV contractility observed with incremental heart rates and so not sufficient to posit a mechanism of the observed reductions in total peripheral resistance beyond increasing cardiac output.

To explore this further, we first examined beat-to-beat BPV, which is a feature of the arterial pressure signal and not dependent on mean arterial pressure or cardiac output. BPV is known to be related to autonomic nervous system activity, and is associated with adverse cardiovascular events following myocardial infarction^23^ and stroke,^24^ and in individuals with hypertension.^25^ We expressed BPV as both the coefficient of variation, regarded as a reliable method which partially corrects for direct proportionality between the average blood pressure and its variation, as well as the ARV, which provides quantification of the absolute average differences between consecutive short term blood pressure readings. Both indices suggested withdrawal of sympathetic tone at the critical heart rate. Given these are indirect proxies derived from the pressure signal, we then examined this directly in a microneurography sub-study in people with HFrEF. We found that with increasing heart rate there was a reduction of single-unit MSNA frequency, which then increased again beyond the peak of the force-frequency relationship curve. Even after accounting for the increased MSNA frequency which would be anticipated at higher heart rates, a reduction in MSNA incidence up to the critical heart rate was observed.

### Physiological and translational implications

Beta-adrenoceptor antagonists are a foundational treatment for HFrEF,^6^ but whether their benefits derive solely from sympathetic antagonism or heart rate lowering *per se* are challenging to disentangle. Our data also suggest that these cannot be separated, with relative unloading of baroreceptors at lower heart rates favouring sympathetic activation, a nadir of sympathetic tone at the peak of the FFR, and increases at higher heart rates as cardiac output falls. These observations may seem at odds with the clinical trial data supporting heart rate lowering strategies, however in SHIFT (Systolic Heart failure treatment with the I*_f_* inhibitor Ivabradine Trial) the greatest benefits of Ivabradine were observed in those with resting heart rates >87bpm, and were not evident in those with heart rates <75bpm who might have been less likely to exceed their critical heart rate during physical activity.^4^ Moreover, in DAVID II (Dual Chamber and VVI Implantable Defibrillator II Trial) a strategy of moderately accelerated atrial-based pacing (AAI-70) did not worsen clinical outcomes compared to back-up ventricular pacing (VVI-40) (*p*=0.95) despite achieving higher mean heart rates at 24 months (73±6bpm vs 65±12bpm at 24 months).^26^

In HFrEF, LV contractility is reduced, the slope of the FFR is less steep, and the critical heart rate is on average lower.^11^ These observations may go some way to explain why approaches to improve exercise tolerance with rate-adaptive pacing have produced largely neutral results.^9,27^ In a prior phase II randomised trial, it was shown that when rate-adaptive pacing algorithms were personalised to not exceed the critical heart rate, exercise distance was longer and LV function was improved at six months.^11^ Reconciling these findings with the present mechanistic data, our findings suggest that the avoidance of both excessively low and excessively high heart rates avoids activation of the sympathetic nervous system, and that the optimal heart rate range is likely to be individual.

## Limitations

This was a prospective, observational study enrolling a representative cohort of participants who had previously received CIEDs for the management of HFrEF. Participants who served as controls had all received CIEDs for the management of bradycardia, and so whilst they did not have HF, these data may not be generalisable to healthy populations. These data also cannot be generalised to physiological changes in heart rate during exercise, where many other factors will change simultaneously. BPV is less well established as a marker of sympathetic tone compared with heart rate variability, and total peripheral resistance was derived as the quotient of mean arterial pressure and cardiac output, however the microneurography data provide independent confirmation that this reflects a mechanistic finding. Finally, our observations are based on acute changes in heart rate, therefore they do not themselves establish a therapeutic strategy or whether increases in heart rate would result in a sustained withdrawal of sympathetic outflow.

## Conclusions

In this prospective, observational, mechanistic study we describe how the FFR relates indices of LV contractility, cardiac output and vascular resistance. Confirming our observations in an exploratory microneurography study, we observed a bidirectional relationship between heart rate and the sympathetic activation, with a nadir of sympathetic tone occurring at the peak of the FFR. Taken together, these data suggest that optimal heart rate targets for HFrEF are likely to be individual.

## Acknowledgements

SS, OIB, MD and RB are supported by the National Institute for Health and Care Research (NIHR). SK, CC and MTK are supported by the British Heart Foundation. MFP and MTK are supported by the NIHR Leeds Biomedical Research Centre. The authors acknowledge the support of the NIHR Leeds Cardiovascular Clinical Research Facility and the NIHR Leeds Biomedical Research Centre.

## Author contributions

JG and KKW conceived the study. SS, CC, BB, JEL, JC and JG collected the data. SS, AG and OIB analysed the data. SS and AG jointly authored the first version of the manuscript. All other authors provided critical revision of the manuscript.

## Data availability statement

Datasets generated and/or analysed from this study are not publicly available due to the inclusion of potentially identifiable information but are available from the corresponding author upon reasonable request.

## Figure legends

**Figure S1.** Representative single-unit action potential from a participant in the microneurography study. <u>Legend</u> Superimposition of single-unit action potentials occurring in diastole confirming similar morphology.

